# Dual hypocretin receptor antagonism enhances sleep and nursing behavior in lactating rats

**DOI:** 10.1101/2025.09.05.674514

**Authors:** Mayda Rivas, Florencia Peña, Clara Prota, Carlos Carrera-Cañas, Miguel Garzón, Pablo Torterolo, Luciana Benedetto

## Abstract

Hypocretins (also known as orexins) are neuropeptides that regulate the sleep-wake cycle and modulate various behaviors, including maternal behavior. They act through two receptor subtypes: hypocretin receptor 1 (HcrtR1) and hypocretin receptor 2 (HcrtR2). Although Dual Orexin Receptor Antagonists (DORAs) are clinically used as hypnotics, most preclinical studies with these drugs have been conducted in males, with limited research in females, leaving the postpartum period largely unexplored. Here, we examined the impact of the DORA Suvorexant on sleep and maternal behavior in lactating rats.

Lactating and virgin female rats were implanted with electrodes for polysomnographic recording. Using a counterbalanced design, Suvorexant was orally administered at doses of 0, 10 (SUV10), and 30 mg/kg (SUV30) to virgin rats in diestrus and to lactating rats between postpartum days 4 and 8. Sleep recordings and maternal behaviors were assessed during the light phase for six hours following the administration of the drug.

Suvorexant reduced wakefulness and increased slow wave sleep, intermediate state, and REM sleep in both groups, with a stronger effect in virgin females. In lactating rats, Suvorexant increased nursing time and milk ejections, while reducing active maternal behavior such as pup-licking.

These findings demonstrate that dual hypocretin receptor antagonism produces hypnotic effects and selectively modulates maternal behavior, promoting nursing while reducing active maternal behavior.

## Introduction

Hypocretins (HCRT), also known as orexins, are two peptides derived from a common pre-pro-HCRT precursor, HCRT-1 (orexin A) and HCRT-2 (orexin B), that exert their biological effects acting via two receptors, HCRT-R1 and HCRT-R2^1,2^. Initially associated with feeding regulation, HCRT are now recognized as key modulators of several functions, such as the promotion of wakefulness and motivated behaviors^3,4^. In addition, it is now well established that a deficiency in HCRT signaling is the underlying cause of narcolepsy ^5–7^.

The postpartum periods induce anatomical and functional changes in the HCRT system. ^8^ reported increased expression of pre-pro-HCRT in rats on postpartum day 1 (PPD1) compared to gestation and PPD14. Similarly, lactating rats show more HCRT-immunoreactive neurons than non-lactating animals ^9–11^. HCRT neuronal activity during lactation, assessed via c-Fos expression, has been shown to increase during postpartum in mice and rats, followed by a decline approaching weaning ^11,12^. Furthermore, increased levels of hypothalamic HCRT-R1 mRNA were found on PPD1 compared to PPD14 and gestation ^8^. These findings suggest that the HCRT system varies along the reproductive stages of the female, likely contributing to the physiological adjustments that occur throughout the different stages of the female’s reproductive cycle. Notably, its activity is heightened in the early postpartum period, which decreases as lactation progresses, supporting its functional relevance during this period ^13^.

Accordingly, previous studies suggest that the HCRT system regulates maternal care. Intermediate intracerebroventricular doses of HCRT-1 enhance grooming of pups, while high doses diminish nursing ^14^. Our prior research demonstrated that microinjection of HCRT-1 into the medial preoptic area (mPOA) promotes active behaviors. In contrast, administration of either a selective HCRT-R1 antagonist or a dual orexin receptor antagonist (DORA) in this area enhances passive behaviors, such as nursing, and increases litter weight gain ^15,16^.

In line with the wake-promoting role of HCRT and its increased activity during the postpartum period, mothers of various species display heightened wakefulness and fragmented sleep during this time ^17–21^.e Indeed, elevated HCRT levels have been associated with poor sleep during pregnancy ^22^, suggesting that this neuropeptide may contribute to the sleep alterations observed after parturition. Consistent with this idea, we previously found that HCRT-1 microinjected into the mPOA increased wakefulness and active maternal behaviors in lactating rats. In contrast, administration of a DORA into the same area improved sleep and nursing, highlighting the role of HCRT in coordinating both processes ^16^.

The wake-promoting function of HCRT has driven the development of DORAs as alternatives to traditional hypnotics ^23,24^. Suvorexant (SUV), the first DORA approved by the U.S. Food and Drug Administration (FDA), fosters a balance between non-REM and REM sleep while preserving overall sleep architecture in individuals with insomnia ^25–27^.

Postpartum sleep disturbances, which are often linked to psychological issues, may benefit from pharmacological interventions ^28^; however, SUV is classified as a category C drug by the FDA for use during pregnancy and lactation due to a lack of comprehensive studies. This underscores the need for further research to clarify the role of HCRT during this critical period of life.

Although we have examined HCRT and DORA actions in the mPOA concerning sleep and maternal behavior, their systemic effects in nursing mothers remain poorly understood. Notably, previous work has shown that postpartum rats exhibit distinct sleep patterns compared to virgins, characterized by increased wakefulness and reduced sleep ^18,19,21^, a pattern considered an adaptive mechanism to support offspring care. Hence, in the present study, we first compared the basal waking and sleep patterns between lactating and virgin females. We then evaluated the systemic effects of SUV on sleep, maternal behavior, and lactation in postpartum rats, and compared the sleep-related outcomes with those observed in virgin females.

## Materials and methods

### Animals and housing

Experiments were conducted in accordance with the Guide for the Care and Use of Laboratory Animals (8th ed., National Academy Press, 2008) and the procedures were approved by the Institutional Animal Care Committee (N° 07151-000022-23). The conditions mirrored those of our previous studies, e.g., ^16,20^; virgin and primiparous Sprague-Dawley female rats (250-300 g) and their pups were used. Pregnant females were housed individually from 2 to 3 days before parturition and remained in this condition until the experiments ended. On PPD1, litters were culled to four female and four male pups per dam. Similarly, virgin females were housed individually throughout the entire experiment. Animals were maintained in a temperature-controlled room (22 ± 1 °C) with a 12-hour light/dark cycle (lights turn on at 6 am) and had unrestricted access to food and water.

### Stereotaxic Surgery

Animals were anesthetized with ketamine, xylazine, and acepromazine (80/2.8/2.0 mg/kg, i.p.), and implanted with nichrome electrodes to record electroencephalogram (EEG) activity from the frontal, parietal, and occipital cortices. A reference electrode was placed in the cerebellum, and a bipolar electrode was positioned in the neck muscles for electromyogram (EMG) recording. The electrodes and connectors were secured to the skull using acrylic cement. After surgery, the animals received sterile saline (0.9%; 10 ml/kg, s.c.), a single dose of ketoprofen (5 mg/kg, s.c.), and a topical antibiotic was applied to the surgical wound.

Surgery for lactating females occurred on PPD1, after which they were returned to their home cages with their pups and housed in a soundproof chamber. Pup growth and development were monitored to confirm that maternal care and lactation were not adversely affected. Similar to lactating rats, virgin females underwent surgery three days before the experiments.

### Experimental Design

Two experimental groups comprising eight animals were used: lactating females (PPD4-8) and virgin females (Figure 1A). Each animal received SUV at doses of 0, 10 (SUV10), or 30 (SUV30) mg/kg, administered in 5 ml/kg of 5% methylcellulose via oral gavage. These doses were based on prior studies demonstrating a standard hypnotic effect in male rats ^25,29,30^. All administrations followed a counterbalanced design with at least one day of rest between experiments (Figure 1A).

**Figure 1.**
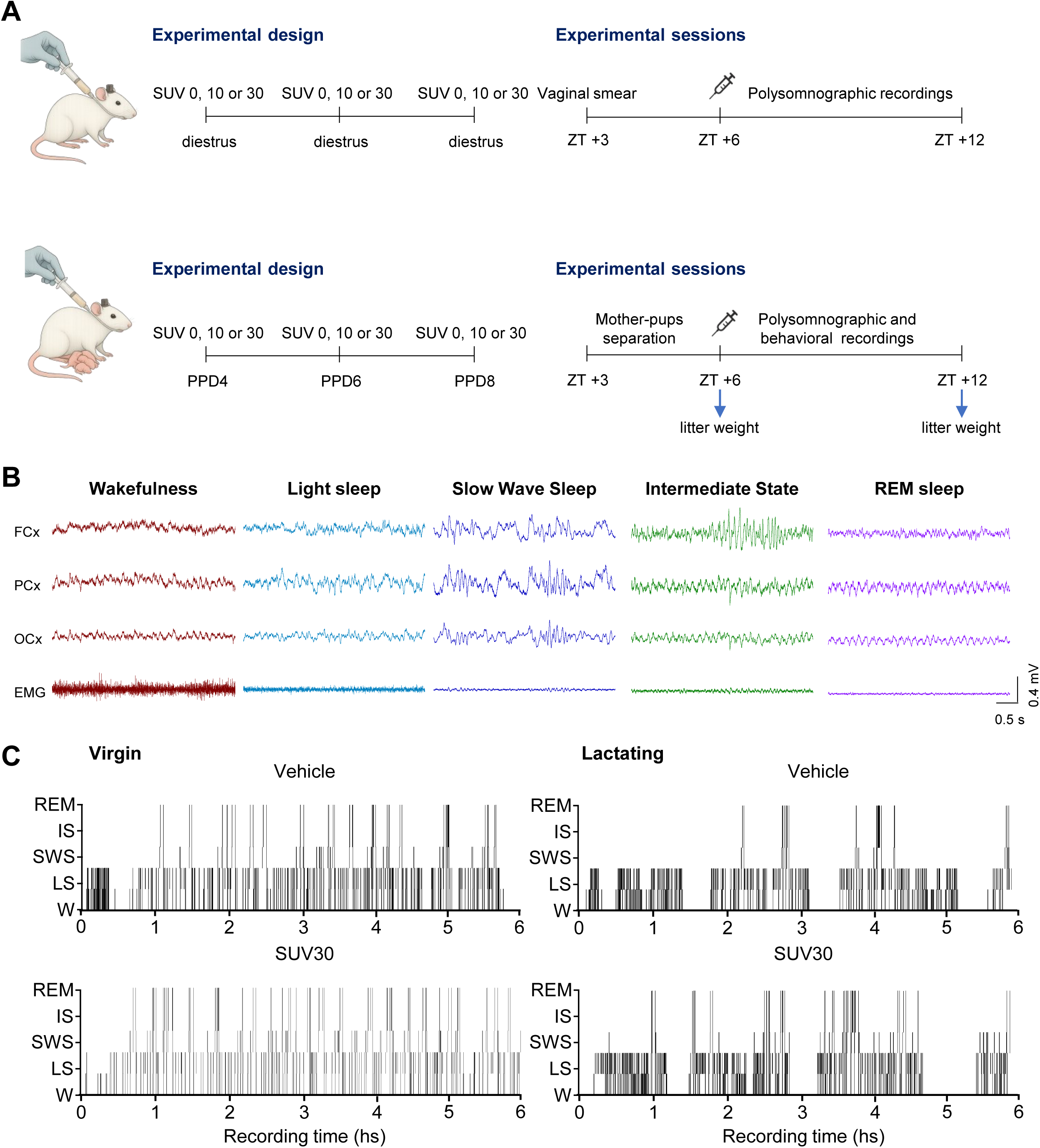
Schematic representation of experimental procedure involving oral administration of Suvorexant (SUV) via gavage in a virgin female (top) and a lactating rat with pups (bottom), both chronically implanted for EEG recordings. (a). Representative EEG traces from frontal (FCx), parietal (PCx), and occipital (OCx) cortices, together with EMG recordings, illustrate the standard criteria for scoring wakefulness and sleep states in rats (b). Representative hypnograms depict baseline sleep architecture in a virgin (left) and a lactating (right) rat following administration of vehicle (top) and SUV30 (bottom) (c). W, wakefulness; LS, light sleep; SWS, slow wave sleep; IS, intermediate state; REM, rapid eye movement sleep.

To assess nursing behavior, pups were separated from their mothers for three hours before the initiation of the drug administration ^31^ (Figure 1A). Five minutes before the end of separation, SUV or vehicle was administered, and pups were weighed. The pups were then returned to the maternal cage, and rats were connected to the polysomnographic system. Polysomnographic and behavioral recordings continued for six hours following the reunion with the pups, from 12:00 to 18:00 (ZT+3 to ZT+12). At the end of the experiments, pups were weighed for a second time to calculate litter weight gain (LWG) (Figure 1A).

In virgin females, SUV was administered, and polysomnographic recordings were conducted similarly to those in lactating females during the same hours of the light phase. The estrous cycle phase correlates with HCRT levels and sleep patterns ^8,32–34^; thus, in virgin rats, vaginal smear samples were collected daily at 9:00 a.m. Samples were dried on glass slides, and estrous stages were identified using standard histological criteria. Experiments were conducted only during the diestrous phase.

At the end of the experiments, the animals were euthanized using an overdose of ketamine and xylazine anesthesia.

### Recording and Analysis of Sleep and Wake Patterns

For polysomnographic recordings, rats were placed in a sound-attenuated, ventilated chamber equipped with independent control of the light/dark cycle and a video camera. They were connected to a slip-ring rotor, allowing continuous recording under freely moving conditions. The bioelectrical signals obtained were amplified (x1000), filtered (low-pass: 100 Hz; high-pass: 10 Hz for EMG and 0.5 Hz for EEG), digitized (512 Hz, 16 bits), and stored for later analysis using Spike2 software. Sleep-wake states were staged in five-second epochs based on standard electrophysiological parameters, as shown in Figure 1B. Wakefulness (W) was characterized by low-amplitude, high-frequency EEG activity and high EMG tone; light sleep (LS) showed intermittent slow waves mixed with desynchronized EEG activity and a moderate reduction in EMG tone; slow-wave sleep (SWS) was identified by the presence of continuous high-amplitude low-frequency EEG waves accompanied by sleep spindles, and low EMG tone; REM sleep was defined by low-amplitude, high-frequency EEG activity and regular theta rhythm in posterior cortical areas, accompanied by muscle atonia. Additionally, the transition state between NREM sleep (LS + SWS) and REM sleep, known as the intermediate state (IS) ^35–37^, was identified by the presence of sleep spindles in the frontal cortex and simultaneous theta activity in posterior cortical areas, along with minimal EMG activity. For this state, three variants were distinguished: entrance-IS, which precedes REM; abortive-IS, which does not precede REM; and exit-IS, which follows the end of REM.

The total time spent in each behavioral state was analyzed across the entire recording period and at hourly intervals. Sleep latencies were measured, defined as the time from recording onset to the first episode of a given state lasting 20 seconds or longer. The number and duration of episodes for each state were also assessed.

### Recording and Analysis of Maternal Behavior

Maternal activity was recorded using a webcam integrated with the polysomnographic acquisition software. The interaction of mothers with their pups was monitored and continuously recorded for six hours. The behaviors were classified into three major categories: hovering over the pups (the dam over the pups while actively engaged in any activity), nursing (being mostly immobile over the pups, characterized by low and high kyphosis and supine postures), and being away from the pups. The duration spent in each behavior was categorized into 5-second epochs and subsequently analyzed in 2-hour windows.

Furthermore, the number of milk ejections was quantified based on the stretching behavior of the pups ^38–40^, and the percentage of LWG was measured as an indirect indicator of milk ejection ^38,41–43^.

Additionally, we analyzed specific active maternal behaviors, including mouthing (rearranging pups within the nest), licking (both anogenital and body licking), and nest building. The analysis was focused on the first two hours following SUV administration, as maternal behavior changes were most pronounced during this period. Due to its low frequency, mouthing was excluded from the analysis.

### Statistics

All values are presented as mean ± S.E.M. (standard error). Data normality was assessed using the Kolmogorov-Smirnov test. Sleep and wake states comparisons were analyzed using a two-way repeated measures ANOVA, with SUV dose as the within-subject factor and physiological condition (virgin or lactating) as the between-subject factor, followed by Tukey’s multiple comparison *post hoc* test when appropriate. Moreover, to assess whether the hypnotic effects of SUV differed between virgin and lactating rats, we calculated the percentage change from vehicle administration for each sleep state following administration of SUV10 and SUV30. These percentage change values were compared between groups using independent-samples t-tests. In addition, effect sizes were estimated using Cohen’s d to quantify the magnitude of the difference between groups.

Maternal behavior states (time in minutes) were analyzed using a one-way repeated measures ANOVA. For non-parametric data, such as time spent in nest-building, the Friedman test was used, followed by Dunn’s *post hoc* test. The criterion used to reject the null hypothesis was *p* < 0.05.

## Results

### Sleep and wake patterns in lactating and virgin rats in baseline conditions

As expected, under baseline conditions, lactating rats spent significantly more time awake and less time in most sleep states (SWS, IS, and REM sleep) compared to virgin females (Table 1). The increased W time in lactating rats was associated with a longer duration of its episodes and occurred in parallel with a reduction in SWS episode duration. Regarding the number of episodes, lactating females exhibited fewer IS episodes as well as fewer REM episodes compared to virgins. Moreover, no differences were observed between lactating and virgin rats in the latency to initiate NREM or REM sleep under baseline conditions (Table 1).

**Table 1.**
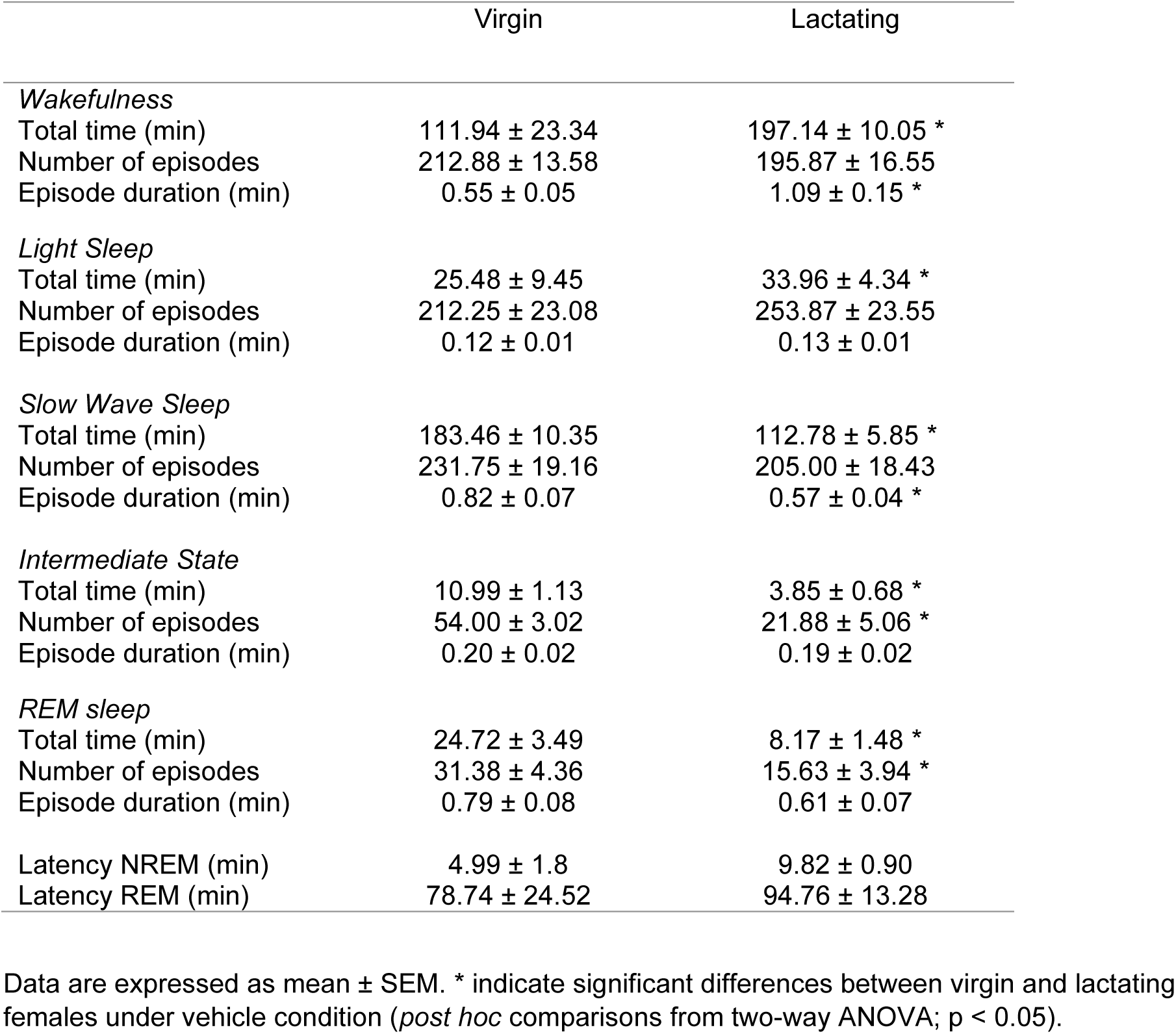
Baseline sleep-wake parameters under vehicle condition in virgin and lactating rats.

### Effect of Suvorexant on sleep and wakefulness

In both virgin and lactating females, SUV administration decreased total W time and increased time spent in SWS, IS, and REM sleep, whereas no changes were observed in LS (Figure 2A, D, G, J, M). As shown in Figure 2B and E, the number of W and LS episodes remained constant following SUV administration in both virgin and lactating females compared to the vehicle. SUV30 increased the number of SWS episodes exclusively in lactating females (Figure 2H), while it enhanced the frequency of IS episodes in both groups (Figure 2K). Significant increases in REM episode frequency were just detected in virgin rats (Figure 2N). Regarding episode duration, only a reduction in W episode length was observed following SUV30 administration in virgin females, whereas no such change occurred in lactating ones (Figure 2C). On the other hand, the latencies to NREM and REM sleep were not significantly affected by SUV administration in either virgin or lactating rats (Figure 2P, Q).

**Figure 2.**
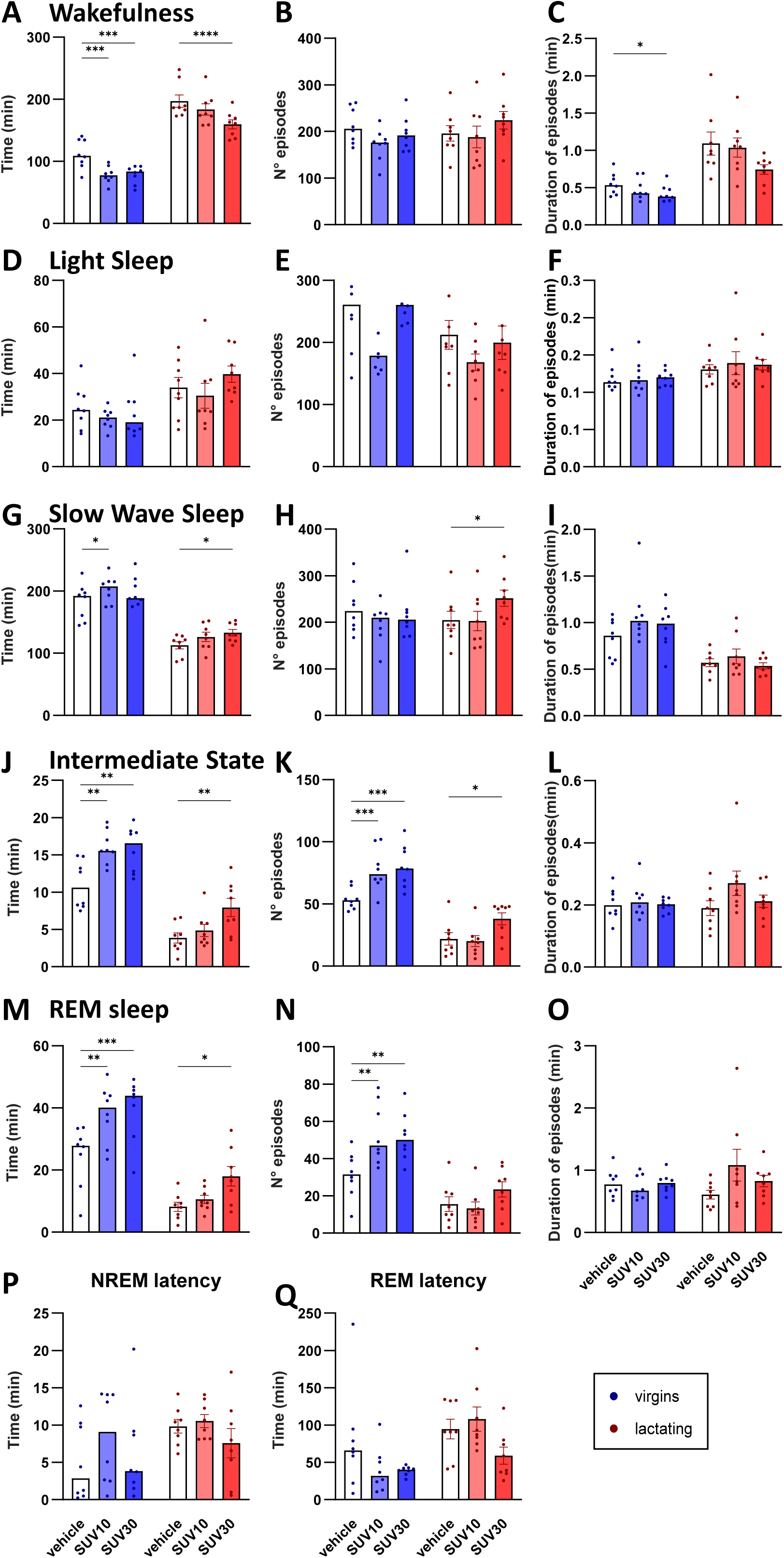
Time spent in wake and sleep states in virgin and lactating rats. (A, D, G, J, M, P); number of episodes (B, E, H, K, N); episode duration (C, F, I, L, O); and latency to NREM (P) and REM sleep (Q) following administration of vehicle, SUV10, and SUV30. Data are presented as mean ± SEM and were analyzed using two-way ANOVA followed by Tukey’s *post hoc* test. \**p* < 0.05, \*\**p* < 0.01, \*\*\**p* < 0.001 indicate significant differences between groups.

Interestingly, the SUV effect was significantly more substantial in virgin females compared to lactating females. A significant interaction between treatment and reproductive condition indicated that SUV produced a greater reduction in total W time in virgin rats compared to lactating rats (SUV × condition F(2, 28) = 3.41, p = 0.047; Figure 2A). Similar interaction effects were observed for the increase in the number of IS episodes (SUV × condition F(2, 28) = 5.05, p = 0.013) and REM episodes (SUV × condition F(2, 28) = 4.95, p = 0.014; Figure 2K and N), suggesting that the impact of SUV varied according to reproductive state, with a stronger effect on virgins. Moreover, we calculated the percentage change from vehicle for total time in each sleep state, and Cohen’s d for the change induced by the administration of SUV10 and SUV30 within each reproductive condition. A significant effect was observed in W time following SUV10, where virgin females exhibited a significantly greater reduction compared to lactating females (virgin: –28.0% ± 6.0, lactating: –6.4% ± 2.7, t(14) = 3.27, p = 0.006; Cohen’s d = 1.63), indicating a larger effect size. Although SUV30 did not reach statistical significance (t(14) = 1.87, p = 0.083), the effect size was large (Cohen’s d = 0.93), suggesting that SUV30 may also reduce W more in virgin than in lactating rats.

Time-course analysis at each 2-hour interval (Figure 3) suggests that the effects of SUV on sleep varied over time. In lactating rats, most changes occurred during the first 2 hours after administration, particularly in the W, SWS, and REM sleep stages (Figures 3B, 3E, and 3I, respectively). In contrast, virgin rats exhibited delayed effects, with significant changes occurring between hours 5 and 6, particularly in W and REM sleep, when both doses had an effect (Figure 3A, 3I). Interestingly, SUV increased IS across the entire recording period in virgin females, whereas in lactating rats, this effect was limited to the 3–4-hour time block.

**Figure 3.**
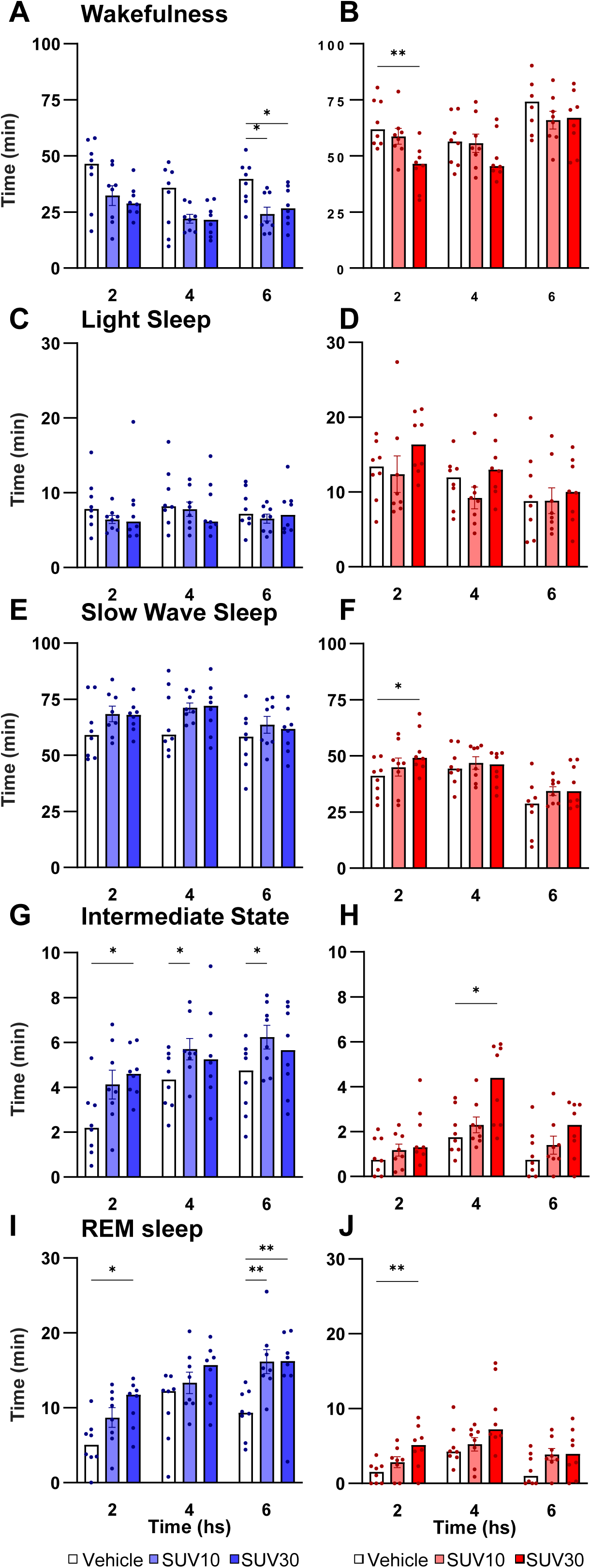
Time spent in wake and sleep states across consecutive two-hour intervals during the 6-h recording period (2: 1–2 h; 4: 3–4 h; 6: 5–6 h) in virgin. (A, C, E, G, I) and lactating (B, D, F, H, J) female rats following administration of vehicle, SUV10, and SUV30. Data are presented as mean ± SEM and were analyzed using one-way ANOVA followed by Tukey’s *post hoc test*. \**p* < 0.05, \*\**p* < 0.01, \*\*\**p* < 0.001 indicate significant differences between SUV doses.

The variations in the different IS subtypes following SUV administration were examined over the total recording time (Figure 4). The administration of both doses of SUV increased the number of Entrance-IS episodes in virgin females, but not in lactating females (Figure 4A). Abortive-IS episodes and exit-IS episodes remained unchanged after SUV administration in both virgin and lactating females (Figure 4B-C).

**Figure 4.**
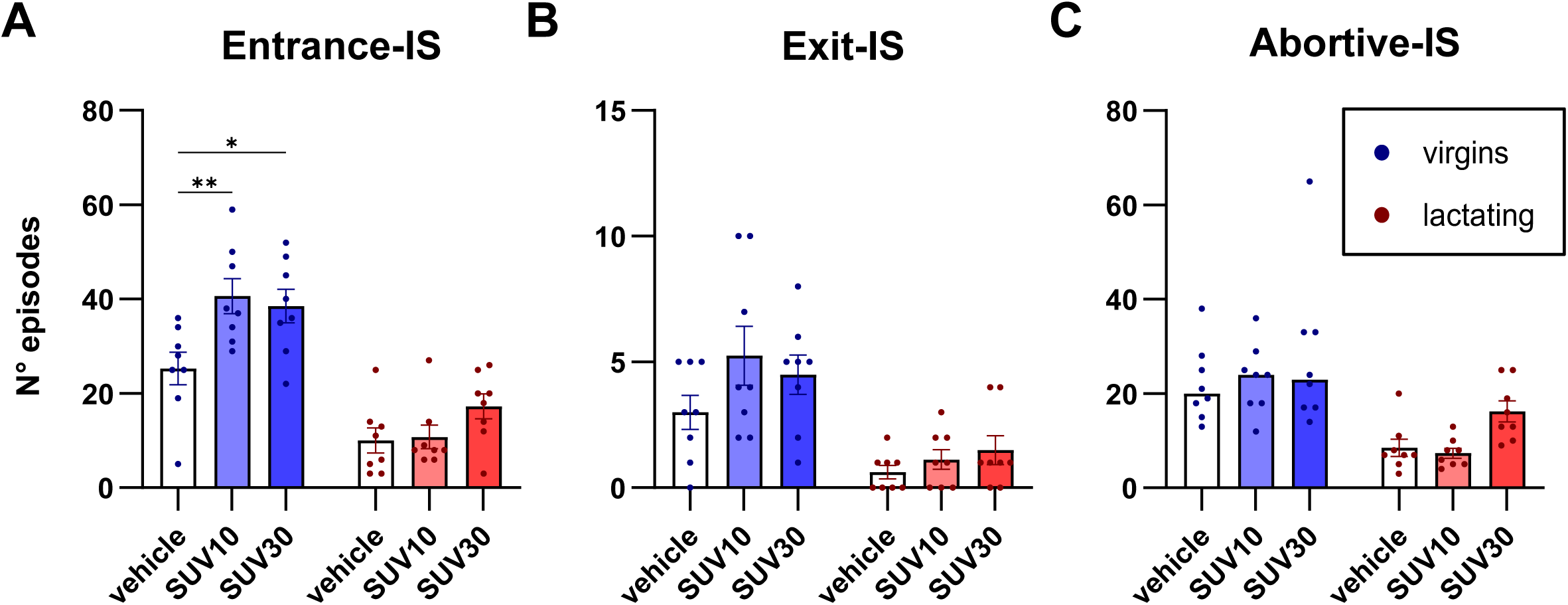
Number of episodes of three IS subtypes: Entrance-IS. (A), Exit-IS (B), and abortive-IS (C) in virgin and lactating rats after administration of vehicle, SUV10, and SUV30. Data are presented as mean ± SEM and were analyzed using two-way ANOVA followed by Tukey’s *post hoc* test, \**p* < 0.05, \*\**p* < 0.01, \*\*\**p* < 0.001 indicate significant differences between groups.

### Maternal behavior

Compared to vehicle administration, both doses of SUV resulted in prolonged nursing time for lactating females (Figure 5A and 5C). This increase was evident in both the total recording time and during the first two hours (Figure 5C and 5F). In addition, SUV30 also increased the number of milk ejections, although it did not affect LWG (Figure 5H and 5I). Additionally, the time spent hovering over the pups decreased following SUV30 administration, while the time spent away from them remained unchanged (Figure 5B and 5D).

**Figure 5.**
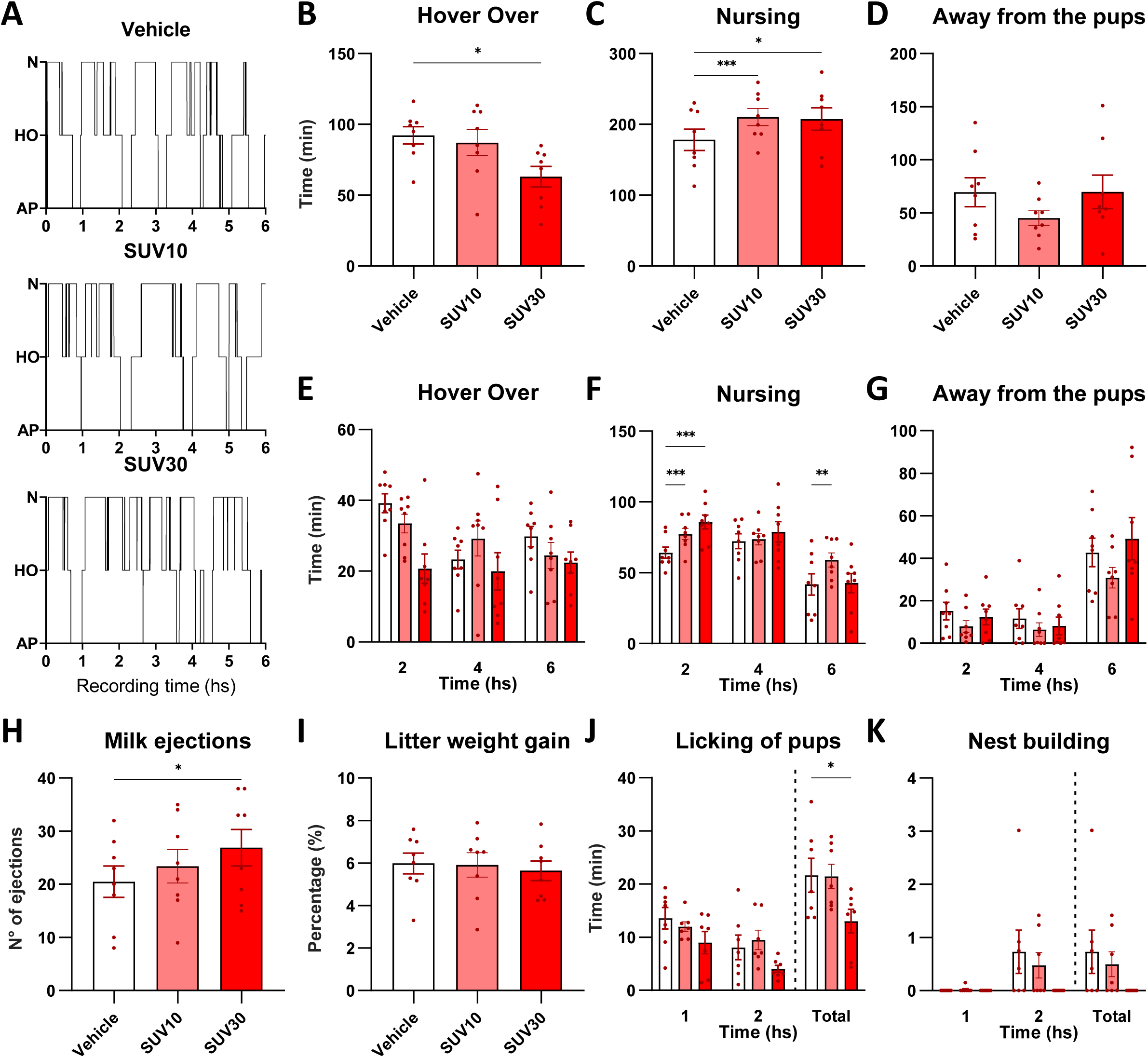
Maternal parameters following administration of vehicle, SUV10, or SUV30. Representative maternograms. (A) based on behavioral staging in 5-s epochs classified into nursing (N), hover over (HO), and away from pups (AP). The Time spent in maternal behaviors during the 6-h recording (B–D) and across consecutive 2-h intervals (2: 1–2 h; 4: 3–4 h; 6: 5–6 h) (E–G). Number of milk ejections (H), percentage of litter weight gain (I), and time spent in active maternal behaviors, including licking of pups (J) and nest building (K). Data are presented as mean ± SEM and were analyzed by one-way ANOVA followed by Tukey’s *post hoc* test, except nest-building, which was analyzed using the Friedman test. \**p* < 0.05, \*\**p* < 0.01, \*\*\**p* < 0.001 indicate significant differences between SUV doses.

Concerning active maternal behavior during the first two hours of recording, SUV30 reduced the time spent licking the pups and tended to reduce nest-building time, although the latter did not reach statistical significance (χ²(2) = 5.44, p = 0.070; Figure 5J and 5K).

## Discussion

This study provides novel insights into the regulation of sleep and maternal behavior during the postpartum period by assessing, for the first time, the effects of the DORA SUV in lactating rats and comparing them with those in virgin rats. Our results indicate that SUV reduced wakefulness and increased sleep in both lactating and virgin rats, exhibiting a more potent wake-inhibitory effect in virgin compared to lactating dams. Furthermore, SUV enhanced nursing behavior and milk ejections without affecting litter weight gain, while diminishing certain active maternal behaviors. The present findings suggest that SUV could potentially serve as an effective hypnotic in mothers without markedly impairing essential maternal behaviors. However, its wake-promoting inhibition appears to be attenuated during lactation. Although further research is still needed to confirm and expand these results, this attenuation may indicate that higher doses could be required to achieve similar hypnotic effects in lactating mothers, a factor that warrants careful evaluation to balance maternal sleep improvement with the preservation of maternal behavior.

### Sleep and wake patterns in lactating and virgin rats

In coherence with previous findings, our data revealed that under control conditions, mother rats spent significantly more time awake and less time in all sleep states, exhibiting longer W episodes, shorter SWS episodes, and fewer IS and REM episodes compared to virgin rats. It is important to note that lactating rats are typically recorded together with their pups (as would be expected in the wild), whereas virgin females are recorded in isolation. In this regard, ^18^ demonstrated that, from postpartum day 2 (PPD2) to PPD20, mother rats exhibited increased W and reduced REM sleep compared to virgin females, based on seven-hour recordings conducted during the light phase, conditions similar to those used in our study. Furthermore, ^19^ reported that postpartum rats showed increased active W during the light phase, with a significant decrease in NREM sleep. However, REM sleep and the frequency of episodes in any state were unaffected. More recently, ^21^ found that postpartum rats at PPD2 display increased W and a reduction in SWS and REM sleep, along with fewer REM episodes during the light phase compared to virgin rats. These earlier studies did not identify IS as a separate sleep state. Here, we demonstrate that IS shows a similar pattern to REM sleep in lactating rats, with both total time and number of episodes reduced in this condition. Despite methodological differences across studies on sleep during lactation, a consistent finding is that W duration increases during the light phase in the early postpartum days, likely serving as an adaptive response to facilitate offspring care. In this sense, previous research in different species has highlighted that sleep patterns during the lactation period are mainly driven by the constant demands of their offspring^44^.

### Effect of Suvorexant on sleep and wake patterns

Our results showed that SUV reduced W while increasing SWS, IS, and REM sleep in lactating and virgin rats, with LS remaining unchanged in both conditions. This hypnotic effect is consistent with preclinical and clinical data that support the sleep-promoting effects of DORAs. Similar to our results, SUV30 administration decreased active W and enhanced NREM and REM sleep for 2 to 7 hours post-administration in male rats ^45,46^. In diverse mammalian species, including rats, dogs, and humans, DORA administered orally during the active period of the circadian cycle decreased alertness and increased NREM and REM sleep ^47^. Recent clinical studies suggest that SUV is clinically safe and effective in improving sleep onset and maintenance, with effects ranging from several weeks to months, in patients with insomnia ^48–51^. Interestingly, pharmacokinetic studies have reported sex-related differences in response to SUV. Female patients showed higher plasma concentrations of SUV than males after the same dose. Women reported somnolence and other adverse effects, and only female participants discontinued driving tests due to excessive drowsiness in safety trials ^49^. However, there is no previous literature about SUV in mother rats. In this regard, we found that SUV effects were more pronounced in virgin rats than in mother rats.

The IS state is the natural transition into REM sleep and is particularly impaired by the absence of HCRT, as observed in narcolepsy, where animals and patients can initiate REM episodes directly from wakefulness, bypassing NREM and IS stages ^52,53^. However, the potential role of HCRT activity in modulating the IS state has not been extensively studied. We thoroughly analyzed IS episodes, categorizing them into three subtypes. The influence of SUV on the number of entrance-IS episodes varied between lactating and virgin females. SUV increased entrance-IS episodes only in virgin females, but did not affect them in lactating females. This increase in the number of entrance-IS in virgin females parallels findings reported in males ^54^. Moreover, these results align with recent findings indicating a pivotal role of HCRT neurons in modulating sleep transitions, since optogenetic inhibition of HCRT neurons in mice has been shown to facilitate the transition from NREM to REM sleep, by increasing the probability and frequency of REM initiation ^55^, and we hypothesize that if measured, it would probably increase IS entrance too. Collectively, these findings support the view that IS is a pivotal transitional state in the sleep–wake cycle, sensitive to modulation by HCRT activity.

To the best of our knowledge, this study is the first to evaluate the hypnotic effects of a drug on postpartum females compared to virgin counterparts. Our results show that SUV, particularly at low doses, reduced total W time more effectively in virgin rats than in lactating ones. Moreover, similar effects were found for the number of IS and REM episodes, with SUV inducing greater increases in virgins compared to lactating females. The increased number, but not duration, of SWS and IS episodes in lactating rats suggests an attempt to recover or consolidate sleep under the influence of SUV. In addition, although SUV increases the number of IS episodes in lactating females, it does not promote the transition from IS to REM, as the number of REM episodes remains unchanged in this group. Besides, the REM facilitation by SUV, although present, is less significant in lactating than in virgin rats. These changes may be caused by the active demands of the pups, which may prevent them from consolidating either SWS, IS, or REM sleep episodes. Finally, abortive-IS and exit-IS episodes remained unchanged after SUV administration in both virgin and lactating females. This could indicate that the effect of HCRT in regulating entry into REM sleep is more important than controlling the intermediate state itself. Thus, our findings contribute to the understanding of how pharmacological HCRT receptor blockade may differentially impact sleep architecture in virgin versus lactating rats, particularly by enhancing REM sleep propensity through modulation of the IS phase in the former group. Further research is needed to elucidate the specific mechanisms by which HCRT modulates IS and its implications in sleep architecture.

The differential response to SUV observed between virgin and lactating females may stem from changes in sleep and W regulation experienced during the postpartum phase, during which various physiological systems adapt to promote W, a critical factor for nurturing offspring. The HCRT system changes during this time could have a significant role ^13^. The reduced ability to decrease W through HCRT receptor blockade in lactating females may be linked to physiological adjustments in the HCRT system that prioritize maternal care. For instance, the postpartum period is associated with increased W and heightened arousal, possibly driven by elevated HCRT tone ^12^. This major activation could counteract the effects of HCRT antagonists, making lactating females less sensitive to the impact of SUV. Alternatively, changes in HCRT receptor expression or sensitivity may contribute to this attenuated response. In support of this hypothesis, ^8^ reported a significant upregulation of HCRT-R1 mRNA and an increased number of HCRT-R1–immunoreactive neurons in the hypothalamus of lactating rats at PPD1. Although these findings do not directly address functional receptor sensitivity, they suggest that the brains of lactating rats may exhibit altered responsiveness to HCRT signaling. Additionally, since HCRT has been shown to modulate prolactin secretion via pituitary receptors ^56^, it is plausible that hormonal states, such as lactation, could influence HCRT system responsiveness. However, to date, no studies have directly examined whether lactation-associated hormonal changes, such as elevated oxytocin or prolactin, affect HCRT receptor function or alter the pharmacological response to HCRT receptor antagonists.

Additionally, the temporal profile of sleep changes induced by SUV differed between virgin and lactating rats. In lactating females, the hypnotic effects were more pronounced during the early post-administration period but dissipated more rapidly than in virgin females, suggesting a shorter efficacy. In this regard, throughout lactation, dynamic alterations in plasma volume, cardiac output, maternal glomerular filtration rate, hematocrit, and serum protein levels can modify both pharmacodynamics and pharmacokinetics ^57^. In male rats, SUV exhibits a terminal half-life of approximately 0.6 hours, with high systemic clearance, and an oral bioavailability of only 20%, indicating rapid elimination following administration {Cox, 2010 #1806, although the sleep effects persist for several hours ^54^. Therefore, the more transient effects observed in lactating rats could reflect a faster systemic elimination of the drug or reduced pharmacodynamic sensitivity in this physiological state. These considerations highlight the significance of reproductive status in influencing both the magnitude and duration of drug effects on sleep–wake regulation.

### Maternal behavior

Our data indicate that SUV increased nursing behavior and the number of milk ejections, while in parallel, it reduced the time the mother spends in active behaviors over the pups, such as hovering over them. In contrast, the time spent away from them remained unchanged. These results are consistent with our previous studies, demonstrating that the administration of DORAs in the mPOA enhances nursing time ^16^. However, in our previous study, nursing increased at the expense of a reduction in the time dams spent away from the pups, without affecting the time spent hovering over them. This difference could imply that the time mothers spend with their pups is less affected by systemic DORA administration than by intra-mPOA, a key area in controlling maternal behavior ^58^.

The blockade of both HCRT receptors enhances nursing duration and the frequency of milk ejections, regardless of whether the drug is administered systemically or directly into the mPOA. As mothers need a resting state to nurse, this effect may be increased indirectly through increased sleep duration, since mother rats often nurse while asleep ^20^. Alternatively, it may suggest the simultaneous facilitation of both nursing and sleep. However, as litter weight gain was not increased by SUV administration, the former hypothesis is more probable.

Additionally, the timing of the drug’s effects varies between oral and local delivery methods. Systemic administration’s peak effect on nursing occurs at two hours, whereas intra-mPOA administration reaches its peak at four hours post-administration. This suggests that blocking the HCRT receptor affects maternal behavior more directly at the systemic level than through the mPOA; it is likely that other regions involved in maternal circuits, such as the prefrontal cortex and the ventral tegmental area, are being impacted by systemic SUV, which may account for the direct effect ^59,60,14^.

Finally, oral administration of SUV reduces active maternal behavior, such as licking the pups. These findings parallel previous results regarding the administration of an antagonist of HCRT-R1 in the mPOA, suggesting that SUV could decrease active maternal care independently of sleep-wake cycle regulation, affecting maternal motivation to care for their offspring. In this context, future studies investigating the effects of DORAs on maternal motivation are critical from a translational perspective, given that DORAs are approved for the treatment of insomnia in several countries ^61^.

### Technical consideration

The method used for drug administration—oral gavage—ensures accurate dosing but is not without limitations. This technique can induce significant stress in experimental animals, potentially affecting physiological and behavioral outcomes, including sleep architecture and maternal behaviors. Moreover, the absorption of orally administered compounds can be influenced by the presence and composition of food in the gastrointestinal tract, introducing inter-individual variability in drug bioavailability and pharmacokinetics. In our study, the drug was prepared as a suspension, a formulation that may result in incomplete dilution and inconsistent homogenization. However, all animals underwent the same procedures, making any significant differences among groups not attributable to this concern.

SUV was administered at mid-light phase to allow evaluation of maternal behavior and comparison with previous findings. However, had it been administered closer to the beginning of the resting phase, its hypnotic effects might have been different— presumably more pronounced—given that HCRT levels are typically elevated during the active phase ^25,62^.

### Conclusions

This study demonstrates that a dual orexin receptor antagonist exerts differential effects on sleep in lactating versus virgin rats, with its hypnotic effects significantly attenuated in lactating females. Notably, in mother rats, HCRT receptor blockade promoted nursing behavior and increased the frequency of milk ejection, without affecting litter weight gain, while reducing active maternal behaviors such as pup licking. Altogether, our results reveal a previously underappreciated interaction between pharmacological sleep promotion and maternal caregiving, highlighting the importance of physiological state in evaluating sleep-inducing treatments. This work contributes to a better understanding of how sleep pharmacotherapies may differentially affect postpartum individuals and highlights potential implications for clinical contexts involving disrupted maternal sleep. Future research is needed to explore further the role of HCRT antagonism in maternal motivation and its impact on postpartum care.

## Bibliography

1. de Lecea L, Kilduff TS, Peyron C, et al. The hypocretins: hypothalamus-specific peptides with neuroexcitatory activity. Proc Natl Acad Sci U S A. 1998; 95 (1): 322–327.

2. Sakurai T, Amemiya A, Ishii M, et al. Orexins and orexin receptors: a family of hypothalamic neuropeptides and G protein-coupled receptors that regulate feeding behavior. Cell. 1998; 92 (5): 1 page following 696.

3. Nishino S, Sakurai T. The orexin/hypocretin system: physiology and pathophysiology. Springer Science & Business Media; 2005.

4. Li SB, de Lecea L. The hypocretin (orexin) system: from a neural circuitry perspective. Neuropharmacology. 2020; 167: 107993.

5. Nishino S, Ripley B, Overeem S, Lammers GJ, Mignot E. Hypocretin (orexin) deficiency in human narcolepsy. Lancet. 2000; 355 (9197): 39–40.

6. Thannickal TC, Moore RY, Nienhuis R, et al. Reduced number of hypocretin neurons in human narcolepsy. Neuron. 2000; 27 (3): 469–474.

7. Seifinejad A, Ramosaj M, Shan L, et al. Epigenetic silencing of selected hypothalamic neuropeptides in narcolepsy with cataplexy. Proc Natl Acad Sci U S A. 2023; 120 (19): e2220911120.

8. Wang JB, Murata T, Narita K, Honda K, Higuchi T. Variation in the expression of orexin and orexin receptors in the rat hypothalamus during the estrous cycle, pregnancy, parturition, and lactation. Endocrine. 2003; 22 (2): 127–134.

9. Sun G, Narita K, Murata T, Honda K, Higuchi T. Orexin-A immunoreactivity and prepro-orexin mRNA expression in hyperphagic rats induced by hypothalamic lesions and lactation. J Neuroendocrinol. 2003; 15 (1): 51–60.

10. Donlin M, Cavanaugh BL, Spagnuolo OS, Yan L, Lonstein JS. Effects of sex and reproductive experience on the number of orexin A-immunoreactive cells in the prairie vole brain. Peptides. 2014; 57: 122–128.

11. Diniz GB, Candido PL, Klein MO, et al. The weaning period promotes alterations in the orexin neuronal population of rats in a suckling-dependent manner. Brain Struct Funct. 2018; 223 (8): 3739–3755.

12. Espana RA, Berridge CW, Gammie SC. Diurnal levels of Fos immunoreactivity are elevated within hypocretin neurons in lactating mice. Peptides. 2004; 25 (11): 1927–1934.

13. Rivas M, Ferreira A, Torterolo P, Benedetto L. Hypocretins, sleep, and maternal behavior. Front Behav Neurosci. 2023; 17: 1184885.

14. D’Anna KL, Gammie SC. Hypocretin-1 dose-dependently modulates maternal behaviour in mice. J Neuroendocrinol. 2006; 18 (8): 553–566.

15. Rivas M, Torterolo P, Ferreira A, Benedetto L. Hypocretinergic system in the medial preoptic area promotes maternal behavior in lactating rats. Peptides. 2016; 81: 9–14.

16. Rivas M, Serantes D, Pena F, et al. Role of Hypocretin in the Medial Preoptic Area in the Regulation of Sleep, Maternal Behavior and Body Temperature of Lactating Rats. Neuroscience. 2021; 475: 148–162.

17. Nishihara K, Horiuchi S. Changes in sleep patterns of young women from late pregnancy to postpartum: relationships to their infants’ movements. Percept Mot Skills. 1998; 87 (3 Pt 1): 1043–1056.

18. Rocha L, Hoshino K. Some aspects of the sleep of lactating rat dams. Sleep Sci. 2009; 2 (2): 88–91.

19. Sivadas N, Radhakrishnan A, Aswathy BS, Kumar VM, Gulia KK. Dynamic changes in sleep pattern during post-partum in normal pregnancy in rat model. Behav Brain Res. 2016; 320: 264–274.

20. Benedetto L, Rivas M, Pereira M, Ferreira A, Torterolo P. A descriptive analysis of sleep and wakefulness states during maternal behaviors in postpartum rats. Arch Ital Biol. 2017; 155 (3): 99–109.

21. Toth A, Petho M, Keseru D, et al. Complete sleep and local field potential analysis regarding estrus cycle, pregnancy, postpartum and post-weaning periods and homeostatic sleep regulation in female rats. Sci Rep. 2020; 10 (1): 8546.

22. Qin L, Luo Y, Chang H, et al. The association between serum orexin-A levels and sleep quality in pregnant women. Sleep Medicine. 2023; 101: 93–98.

23. Yang LP. Suvorexant: first global approval. Drugs. 2014; 74 (15): 1817–1822.

24. Scott LJ. Lemborexant: First Approval. Drugs. 2020; 80 (4): 425–432.

25. Gotter AL, Garson SL, Stevens J, et al. Differential sleep-promoting effects of dual orexin receptor antagonists and GABAA receptor modulators. BMC Neurosci. 2014; 15: 109.

26. Snyder E, Ma J, Svetnik V, et al. Effects of suvorexant on sleep architecture and power spectral profile in patients with insomnia: analysis of pooled phase 3 data. Sleep Med. 2016; 19: 93–100.

27. Monti JM, Torterolo P, Pandi-Perumal SR. The Effects of Benzodiazepine and Nonbenzodiazepine Agents, Ramelteon, Low-dose Doxepin, Suvorexant, and Selective Serotonin 5-HT2A Receptor Antagonists andmInverse Agonists on Sleep and Wakefulness. Clinical Medicine Insights: Therapeutics. 2016; 8: 29–36.

28. Palagini L, Bramante A, Baglioni C, et al. Insomnia evaluation and treatment during peripartum: a joint position paper from the European Insomnia Network task force “Sleep and Women,” the Italian Marce Society and international experts task force for perinatal mental health. Arch Womens Ment Health. 2022; 25 (3): 561–575.

29. Sanchez-Alavez M, Benedict J, Wills DN, Ehlers CL. Effect of suvorexant on event-related oscillations and EEG sleep in rats exposed to chronic intermittent ethanol vapor and protracted withdrawal. Sleep. 2019; 42 (4).

30. Gamble MC, Katsuki F, McCoy JG, Strecker RE, McKenna JT. The Dual Orexin Receptor Antagonist DORA-22 Improves Mild Stress-induced Sleep Disruption During the Natural Sleep Phase of Nocturnal Rats. Neuroscience. 2021; 463: 30–44.

31. Stern JM, Lonstein JS. Neural mediation of nursing and related maternal behaviors. Prog Brain Res. 2001; 133: 263–278.

32. Porkka-Heiskanen T, Kalinchuk A, Alanko L, Huhtaniemi I, Stenberg D. Orexin A and B levels in the hypothalamus of female rats: the effects of the estrous cycle and age. Eur J Endocrinol. 2004; 150 (5): 737–742.

33. Silveyra P, Catalano PN, Lux-Lantos V, Libertun C. Impact of proestrous milieu on expression of orexin receptors and prepro-orexin in rat hypothalamus and hypophysis: actions of Cetrorelix and Nembutal. Am J Physiol Endocrinol Metab. 2007; 292 (3): E820–828.

34. Toth A, Keseru D, Petho M, et al. Sleep and local field potential effect of the D2 receptor agonist bromocriptine during the estrus cycle and postpartum period in female rats. Pharmacol Biochem Behav. 2024; 239: 173754.

35. Sánchez-López AS-P M.; Escudero, M. Temporal dynamics of the transition period between nonrapid eye movement and rapid eye movement sleep in the rat. Sleep. 2018; 41 (9).

36. Carrera-Canas C, Garzon M, de Andres I. The Transition Between Slow-Wave Sleep and REM Sleep Constitutes an Independent Sleep Stage Organized by Cholinergic Mechanisms in the Rostrodorsal Pontine Tegmentum. Front Neurosci. 2019; 13: 748.

37. Serantes D, Cavelli M, Gonzalez J, Mondino A, Benedetto L, Torterolo P. Characterising the power spectrum dynamics of the non-REM to REM sleep transition. J Sleep Res. 2024: e14388.

38. Lincoln DW, Hill A, Wakerley JB. The milk-ejection reflex of the rat: an intermittent function not abolished by surgical levels of anaesthesia. J Endocrinol. 1973; 57 (3): 459–476.

39. Voloschin LM, Tramezzani JH. Milk ejection reflex linked to slow wave sleep in nursing rats. Endocrinology. 1979; 105 (5): 1202–1207.

40. Peña F, Rivas M, Gonzalez J, et al. Sleep and maternal behavior in the postpartum rat after haloperidol and midazolam treatments. Sleep Science. 2020; 13 (Special Issue): 78–86.

41. Peña F, Rivas M, Serantes D, Ferreira A, Torterolo P, Benedetto L. Is sleep critical for lactation in rat? Physiol Behav. 2023; 258: 114011.

42. Peña F, Serantes D, Rivas M, et al. Acute and chronic sleep restriction differentially modify maternal behavior and milk macronutrient composition in the postpartum rat. Physiol Behav. 2024; 278: 114522.

43. Stern JM, Taylor LA. Haloperidol inhibits maternal retrieval and licking, but enhances nursing behavior and litter weight gains in lactating rats. J Neuroendocrinol. 1991; 3 (6): 591–596.

44. Benedetto L, Pena F, Rivas M, Ferreira A, Torterolo P. The Integration of the Maternal Care with Sleep During the Postpartum Period. Sleep Med Clin. 2023; 18 (4): 499–509.

45. Cox CD, Breslin MJ, Whitman DB, et al. Discovery of the dual orexin receptor antagonist [(7R)-4-(5-chloro-1,3-benzoxazol-2-yl)-7-methyl-1,4-diazepan-1-yl][5-methyl-2-(2H-1,2,3-triazol-2-yl)phenyl]methanone (MK-4305) for the treatment of insomnia. J Med Chem. 2010; 53 (14): 5320–5332.

46. Winrow CJ, Gotter AL, Cox CD, et al. Promotion of sleep by suvorexant-a novel dual orexin receptor antagonist. J Neurogenet. 2011; 25 (1-2): 52–61.

47. Brisbare-Roch C, Dingemanse J, Koberstein R, et al. Promotion of sleep by targeting the orexin system in rats, dogs and humans. Nature medicine. 2007; 13 (2): 150–155.

48. Herring WJ, Connor KM, Ivgy-May N, et al. Suvorexant in Patients With Insomnia: Results From Two 3-Month Randomized Controlled Clinical Trials. Biol Psychiatry. 2016; 79 (2): 136–148.

49. Norman JL, Anderson SL. Novel class of medications, orexin receptor antagonists, in the treatment of insomnia – critical appraisal of suvorexant. Nat Sci Sleep. 2016; 8: 239–247.

50. Han AH, Burroughs CR, Falgoust EP, et al. Suvorexant, a Novel Dual Orexin Receptor Antagonist, for the Management of Insomnia. Health Psychol Res. 2022; 10 (5): 67898.

51. Khazaie H, Sadeghi M, Khazaie S, Hirshkowitz M, Sharafkhaneh A. Dual orexin receptor antagonists for treatment of insomnia: A systematic review and meta-analysis on randomized, double-blind, placebo-controlled trials of suvorexant and lemborexant. Front Psychiatry. 2022; 13: 1070522.

52. Schoch SF, Werth E, Poryazova R, Scammell TE, Baumann CR, Imbach LL. Dysregulation of Sleep Behavioral States in Narcolepsy. Sleep. 2017; 40 (12).

53. Chemelli RM, Willie JT, Sinton CM, et al. Narcolepsy in orexin knockout mice: molecular genetics of sleep regulation. Cell. 1999; 98 (4): 437–451.

54. Carrera-Canas C, De Andres IT, Callejo M, Garzon M. Hypocretinergic/Orexinergic transmission: volume versus synaptic neuropeptides release. Differential roles of the hypocretinergic/orexinergic receptors. In: proceedings from the JOURNAL OF SLEEP RESEARCH; 2024.

55. Ito H, Fukatsu N, Rahaman SM, et al. Deficiency of orexin signaling during sleep is involved in abnormal REM sleep architecture in narcolepsy. Proceedings of the National Academy of Sciences. 2023; 120 (41): e2301951120.

56. Molik E, Zieba DA, Misztal T, et al. The role of orexin A in the control of prolactin and growth hormone secretions in sheep--in vitro study. J Physiol Pharmacol. 2008; 59 Suppl 9: 91–100.

57. Deferm N, Dinh J, Pansari A, Jamei M, Abduljalil K. Postpartum changes in maternal physiology and milk composition: a comprehensive database for developing lactation physiologically-based pharmacokinetic models. Frontiers in Pharmacology. 2025; 16: 1517069.

58. Numan M. Medial preoptic area and maternal behavior in the female rat. J Comp Physiol Psychol. 1974; 87 (4): 746–759.

59. Kallo I, Omrani A, Meye FJ, de Jong H, Liposits Z, Adan RAH. Characterization of orexin input to dopamine neurons of the ventral tegmental area projecting to the medial prefrontal cortex and shell of nucleus accumbens. Brain Struct Funct. 2022; 227 (3): 1083–1098.

60. Vittoz NM, Berridge CW. Hypocretin/orexin selectively increases dopamine efflux within the prefrontal cortex: involvement of the ventral tegmental area. Neuropsychopharmacology. 2006; 31 (2): 384–395.

61. Sateia MJ, Buysse DJ, Krystal AD, Neubauer DN, Heald JL. Clinical Practice Guideline for the Pharmacologic Treatment of Chronic Insomnia in Adults: An American Academy of Sleep Medicine Clinical Practice Guideline. J Clin Sleep Med. 2017; 13 (2): 307–349.

62. Taheri S, Sunter D, Dakin C, et al. Diurnal variation in orexin A immunoreactivity and prepro-orexin mRNA in the rat central nervous system. Neurosci Lett. 2000; 279 (2): 109–112.

